# Analysis of nasopharyngeal microbiome patterns in Zambian infants with fatal acute febrile illness

**DOI:** 10.1101/2023.09.27.559805

**Authors:** Aubrey R. Odom, Jessica McClintock, Christopher J. Gill, Rachel Pieciak, Arshad Ismail, William B. MacLeod, W. Evan Johnson, Rotem Lapidot

## Abstract

**Introduction:** Associative connections have previously been identified between nasopharyngeal infections and infant mortality. The nasopharyngeal microbiome may potentially influence the severity of these infections.

**Methods:** We conducted an analysis of a longitudinal prospective cohort study of 1,981 infants who underwent nasopharyngeal sampling from 1 week through 14 weeks of age at 2–3-week intervals. In all, 27 microbiome samples from 9 of the infants in the cohort who developed fatal acute febrile illness (fAFI) were analyzed in pooled comparisons with 69 samples from 10 healthy comparator infants. We completed 16S rRNA amplicon gene sequencing all infant NP samples and characterized the maturation of the infant NP microbiome among the fAFI(+) and fAFI(-) infant cohorts.

**Results:** Beta diversity measures of fAFI(-) infants were markedly higher than those of fAFI(+) infants. The fAFI(+) infant NP microbiome was marked by higher abundances of *Escherichia, Pseudomonas, Leuconostoc*, and *Weissella*, with low relative presence of *Alkalibacterium, Dolosigranulum, Moraxella*, and *Streptococcus*.

**Conclusions:** Our results suggest that nasopharyngeal microbiome dysbiosis precedes fAFI in young infants. Early dysbiosis, involving microbes such as *Escherichia*, may play a role in the causal pathway leading to fAFI or could be a marker of other pathogenic forces that directly lead to fAFI.

## Introduction

Fatal acute febrile illness (fAFI) is a common end point for many infectious diseases. In children under five years old in low-middle income countries (LMICs), infectious diseases remain a leading cause of death, including respiratory infections, meningitis, sepsis, and diarrhea^1,2^. For many pathogens that commonly cause infections in this age group (e.g., *Streptococcus pneumoniae, Haemophilus influenzae* and *Neisseria meningitidis*), the nasopharynx is the ecological niche for early colonization and entry into the host. The nasopharyngeal (NP) microbiome, and the complex interactions between commensals and pathogenic microorganisms, can affect health by providing ‘colonization resistance’ to protect against pathogen invasion by influencing the environment in the nasopharynx (pH, metabolites, excretion of antibiotics etc.) and by triggering the immune system^3,4^. A range of respiratory diseases have been linked to aberrations in the NP microbiome, including pneumonia^5^, RSV bronchiolitis^6-8^, general LRTIs^9^, otitis media^10-13^, and asthma^14^.

In this analysis, we investigated nasopharyngeal microbiome changes in a small longitudinal cohort of Zambian infants with fatal acute febrile illness (fAFI). We addressed the following questions: 1) how does the longitudinal NP microbiome of infants who died of fAFI differ from healthy comparator infants; 2) are there specific identifiable patterns in those infants leading up to and at the time of fAFI; and 3) how do we interpret these findings in terms of their associative relationship with fAFI? We hypothesized that changes in the microbiome leading up to the infection itself could be observed early in life, before fAFI. Our findings suggest that NP microbiome dysbiosis precedes fAFI, sometimes before the onset of initial infection.

## Methods

### Study Design and Population

We identified study infants from an existing longitudinal NP sample library collected in Lusaka, Zambia between 2015 and 2016. The sample library was part of a nested time-series case comparator study within the prospective longitudinal Southern Africa Mother-Infant Pertussis study (SAMIPS)^15^. In that study, infants (and their mothers) underwent NP sampling from after birth through 14 weeks of age at 2–3-week intervals, while capturing contemporaneous clinical data, including illnesses, exposure to antibiotics, and routine vaccines received. The institutional review boards at Boston Medical Center and Excellence in Research Ethics and Science Converge in Lusaka jointly provided ethical oversight (The ERES Converge, Lusaka. REF# 2015-Jan-002, Date: 01/02/2015; BUMC IRB, Boston. # H-33521, Date: 12/12/2014). All mothers provided written informed consent, with consent forms presented in English and the two dominant vernacular languages spoken in Lusaka: Bemba and Nyanja. All infants enrolled in SAMIPS were born at term (>37 weeks), were not underweight (>2500 grams), had no acute or chronic conditions known at the time of enrollment, were not born via cesarean section, and had no known complications during pregnancy or labor and delivery.

We chose a subset of nine case infants with fAFI and ten healthy comparator infants, comprising 96 NP swabs. The selection criteria are as follows: The ten fAFI(-) infant samples were healthy controls sourced from a previous study^9^ on infant LRTI development^16^. The fAFI(+) cohort includes infants who died of sepsis during the study period as indicated in the severe adverse event (SAE) forms. Our team completed adjudication of the deidentified SAE forms and only included infants who died from an infectious cause, defined by fever with additional symptoms like a cough, difficulty breathing, diarrhea, or poor feeding. Nine infants met these criteria for fAFI. Insufficient information was available to classify deaths as sepsis induced, although symptoms align with this diagnosis. All infants that were diagnosed with fAFI were given an antibiotic (often classified in the SAEs as a treatment for sepsis).

Characteristics of the study cohort are delineated in **Table 1**. Some missingness in the data is present as the fAFI infants have fewer than seven collected time points having died before the end of the study. Of the nine infants, the fAFI group was subdivided into two groups with final samples at time point 1 (n=4) and time points 3_LJ_4 (n=5). For all infants, sepsis onset was within 12 days of the last or penultimate sample (Figure 1). fAFI(-) comparator infants were selected to have complete sample collections at all scheduled time points.

**Table 1:**
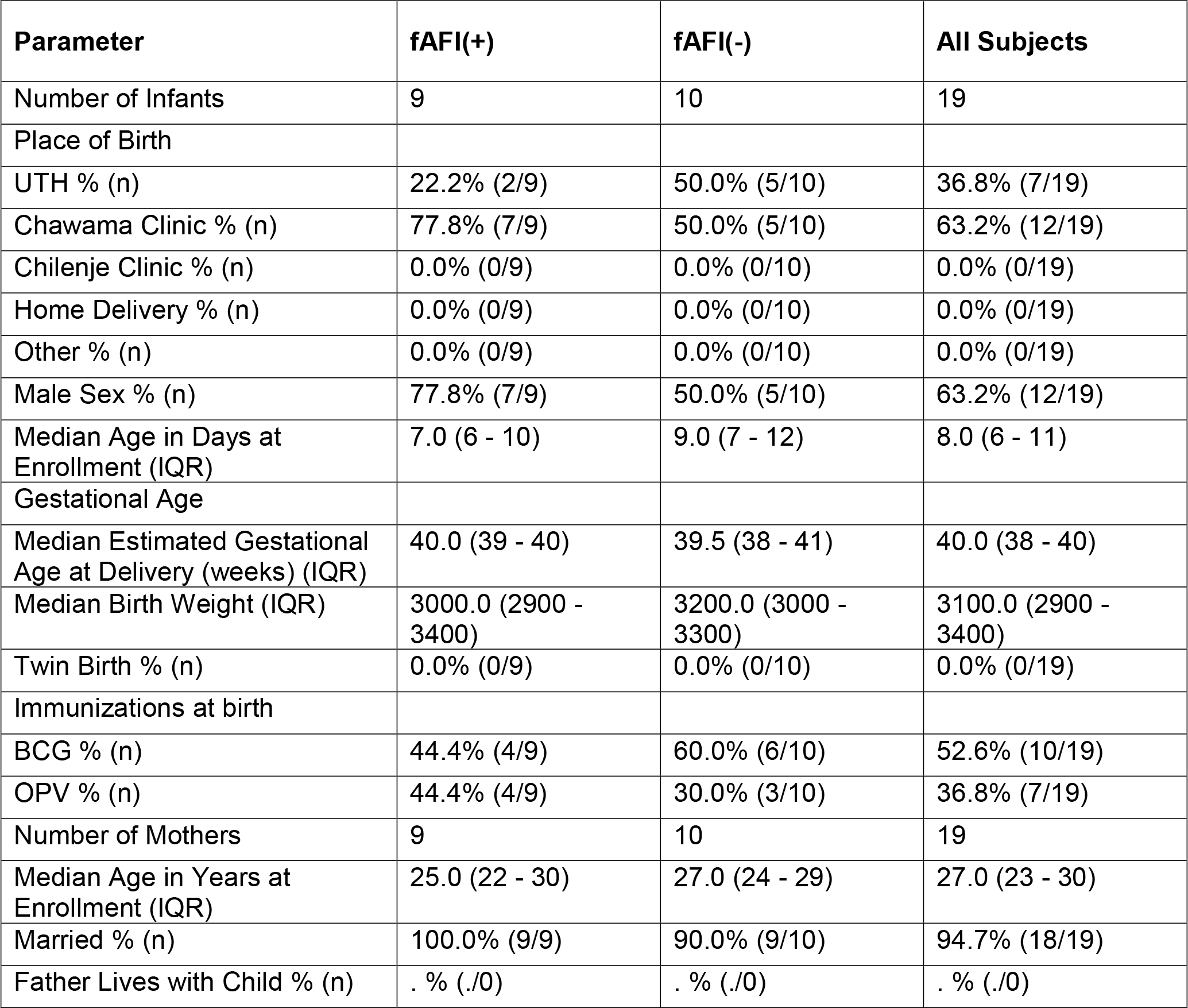
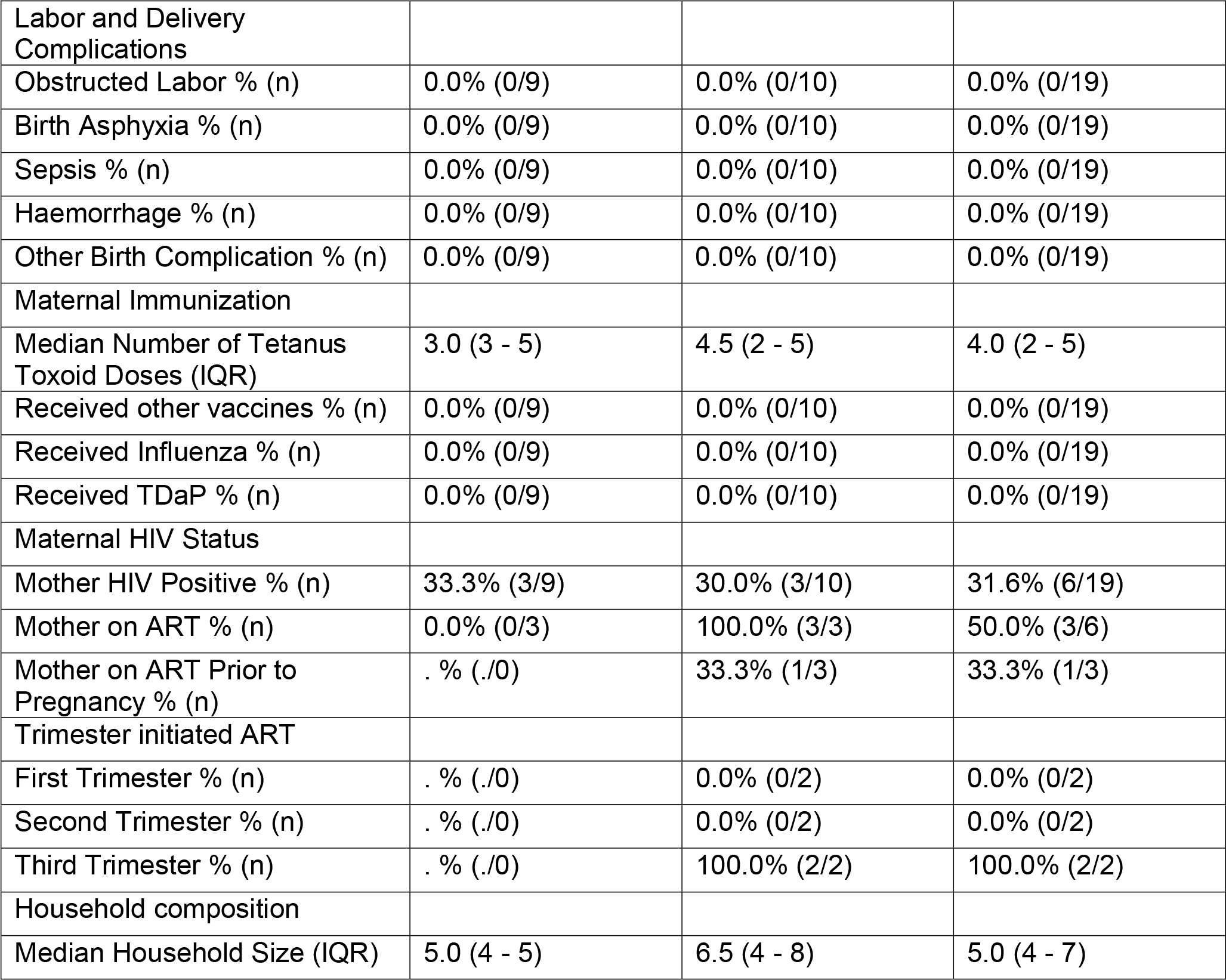
Baseline demographic characteristics of the full infant cohort, stratified by fatal acute febrile illness (fAFI) status. UTH refers to whether the infants was born in the University Teaching Hospital, IQR refers to the interquartile range, BCG refers to the Bacille Calmette-Guérin vaccine for tuberculosis disease, and OPV refers to oral polio vaccine. ART refers to antiretroviral therapy.

**Figure 1:**
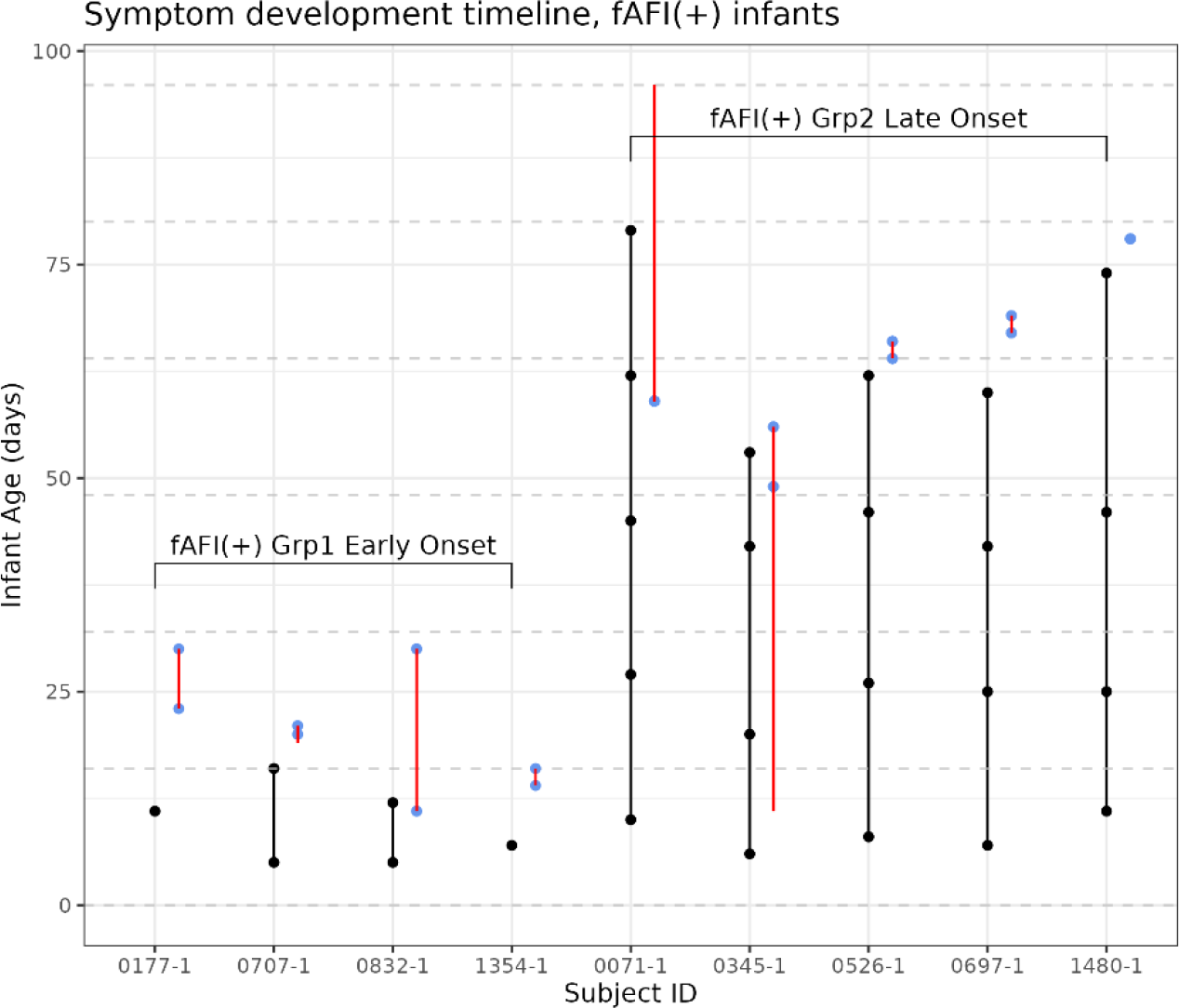
A vertical line plot illustrating the number of samples available for each of the infants with fatal acute febrile illness (fAFI) and the infant age at which they were taken. Red lines indicate periods when the infants had been reported as having antecedent sickness of any kind. Blue dots indicate the start/end of fAFI symptoms. Black dots indicate NP sampling visits for the study, connected by a black line. Samples are separated based on early or late onset, demarcated by the early and late group respectively.

Details on sample collection and processing have been made available as part of the supplementary materials.

### Statistical analysis

All analyses were performed using R Statistical Software (v4.2.1; R Core Team 2022).

The R package ggplot2 v3.4.0 was used to visualize the relative abundance of genera across sample groups with stacked bar plots and spaghetti plots. For stacked bar plots, samples were combined within a given time point and fAFI status group and plotted as a single bar. Spaghetti plots were constructed separately for each genus, split according to fAFI status, and colored by early or late fAFI onset, denoted as fAFI(+) groups 1 and 2, respectively. Genus relative abundances in log CPM were plotted over time.

A uniform manifold approximation and projection UMAP analysis was used to group all infant samples, with nearest neighbors set to 10 and using a Manhattan distance metric.^17^ The UMAP plot was colored according to three separate classifications; time point, subject ID, and fAFI status.

We computed the Bray-Curtis dissimilarity index using the animalcules^18^ v1.9.3 and vegan^19^ v2.6.4 packages. Using the dissimilarity matrix, we performed non-metric multidimensional scaling (NMDS) with the metaMDS function from the vegan^19^ package to localize samples in two dimensional ordination space for a maximum of 500 iterations. We utilized a stress score upper bound of 0.2 as a cutoff for an acceptable reduced-data representation of the data.^20^ The function envfit was then used to perform a multiple regression and permutation test on the variables of time point and fAFI status. The calculated R2 value represents the variation explained by the model of multiple regression, and the resulting p-value represents the goodness of fit of the variable in relation to the ordination axes, with the null hypothesis that there is no relation. Packages ggplot2 v3.4.0 and ggforce v0.4.1 were used to generate NMDS plots with ellipses encircling the data alongside centroids of the fAFI(-) and fAFI(+) subgroups.

Given the small number of samples available between early and late fAFI(+) subgroups, we relied on the two-sample Student’s t-test to compare normalized log CPM genus abundances across groups and time points. This method is appropriate for small sample sizes as it is robust to the assumption of normality and can provide reliable inference in the face of increased uncertainty. We performed five separate tests. We compared the early fAFI(+) group to the fAFI(-) group at time point 1, then compared fAFI(+) late onset group to the fAFI(-) at each of the time points 1–3, and then combined at 4/5. If an infant had more than 2 samples within a timeframe, only the sample closest to fAFI onset was used. At each pairing, we conducted nested tests for all genera and “Other” microbes, then corrected p-values using the Bonferroni correction. We then separately conducted tests for all genera between fAFI(+) infants only. For each genus, we performed a two-sample t-test between fAFI(+) early onset infants at time point 1 versus late onset infants combined at time points 3 and 4, again limiting each infant to the sample closest to AFI onset period.

## Results

### Sequencing

Our FastQC analysis indicated that the overall sequencing quality was excellent for the fAFI(-) infants, which were sequenced in a different run from the AFI(+) infants. For the former, mean Phred quality scores were greater than 25 (99.5% accuracy) from the 25 to 225 bp region for both forward and reverse reads. For the fAFI(+) infants, Phred quality scores were greater than 25 (99.5% accuracy) from the 25 to 200 bp region for both forward and reverse reads. Trimmomatic dropped less than 3% of reads in any given sample, with around 10% surviving with only forward or reverse reads.

The downstream analysis covered 96 NP swab samples, with an average of 103,181 ± 15,447 (mean ± SE) reads per sample (min = 26,297; max = 1,475,543). In our raw data, we uniquely identified 34 phyla, 1530 genera, and 3472 total species across all samples.

### Relative Abundance

We generated stacked bar plots to compare the relative abundance of microbes by timepoint across cohorts (**Figure 2**). The fAFI(-) infants have the same genera consistently appearing across timepoints in contrast to the fAFI(+) groups (Figure 2). In fAFT(-) infants at timepoint 1, we observed a predominance of skin bacteria, notably *Staphylococcus* and *Corynebacterium*, followed by the emergence of and replacement by more typical respiratory bacteria, notably *Streptococcus, Haemophilus*, and *Moraxella* as the infants aged. *Dolosigranulum* was detected at all time points. By comparison, in the fAFI(+) infants, *Leuconostoc, Novosphingobium, Weissella*, and *Escherichia* are all elevated in the microbial profiles across timepoints. The fAFI(+) infants are also distinguished by the frequent presence of *Pseudomonas* and *Escherichia* and the complete absence of *Dolosigranulum*.

**Figure 2:**
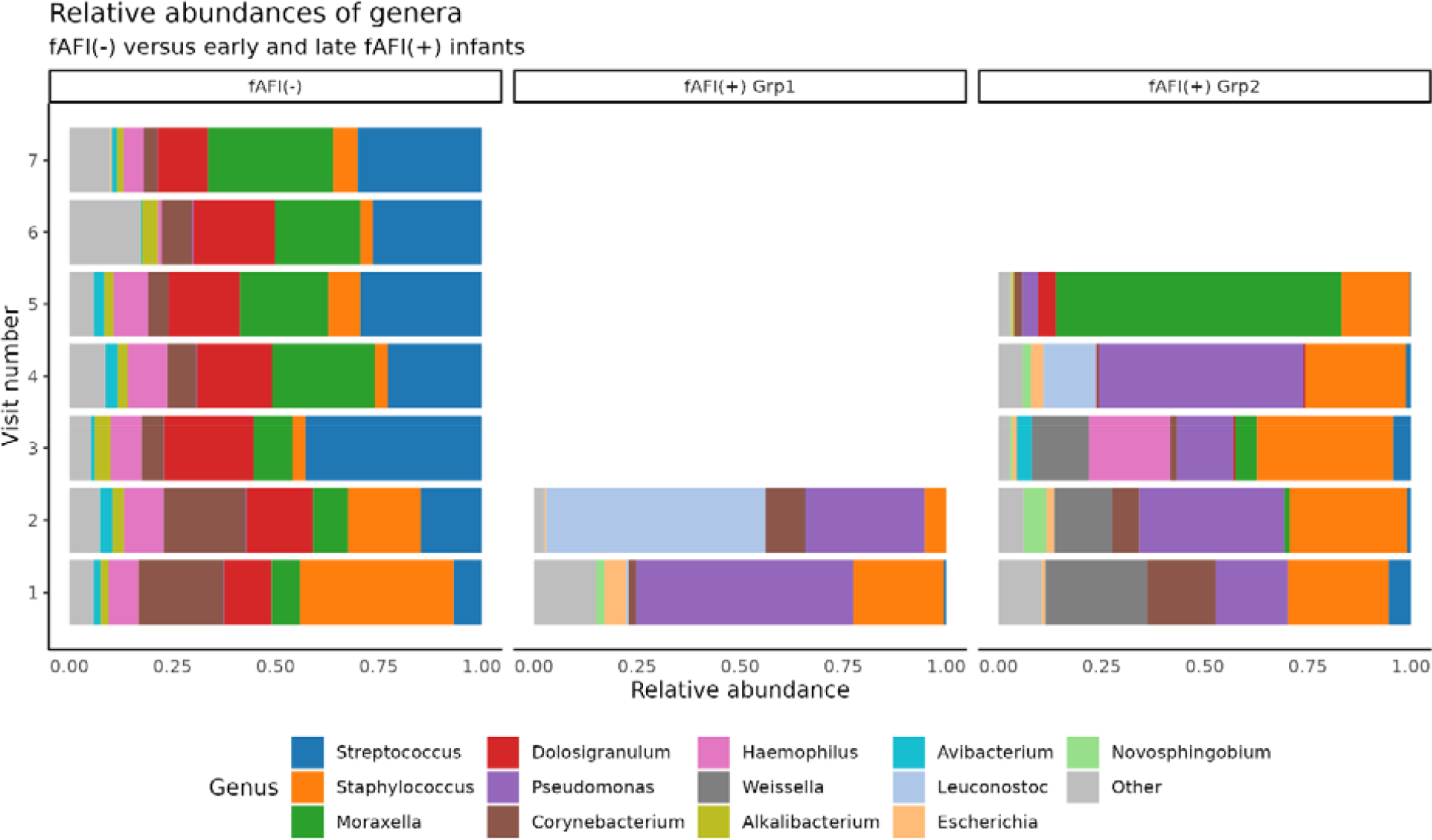
A stacked bar plot comparing the microbial genera profiles of infants with fatal acute febrile illness (fAFI) in each of two different groups (group 1 early onset and group 2 late onset) with control fAFI(-) infants. Note the large discrepancies in the abundance of certain genera in each group, including the relative lack of *Dolosigranulum* (red) and increased relative abundance of *Escherichia* (light orange), *Pseudomonas* (purple), and *Leuconostoc* (blue) in fAFI(+) infants.

We then created spaghetti plots for each genus to provide a linearly distinct view of differences between samples and groups, highlighting similarities in the fAFI(+) groups (Figure 3). *Streptococcus* and *Alkalibacterium* are in decreased abundance across all timepoints in the AFI(+) infants, whereas the *Escherichia* and *Pseudomonas* genera are higher in abundance in fAFI(+) infants.

**Figure 3:**
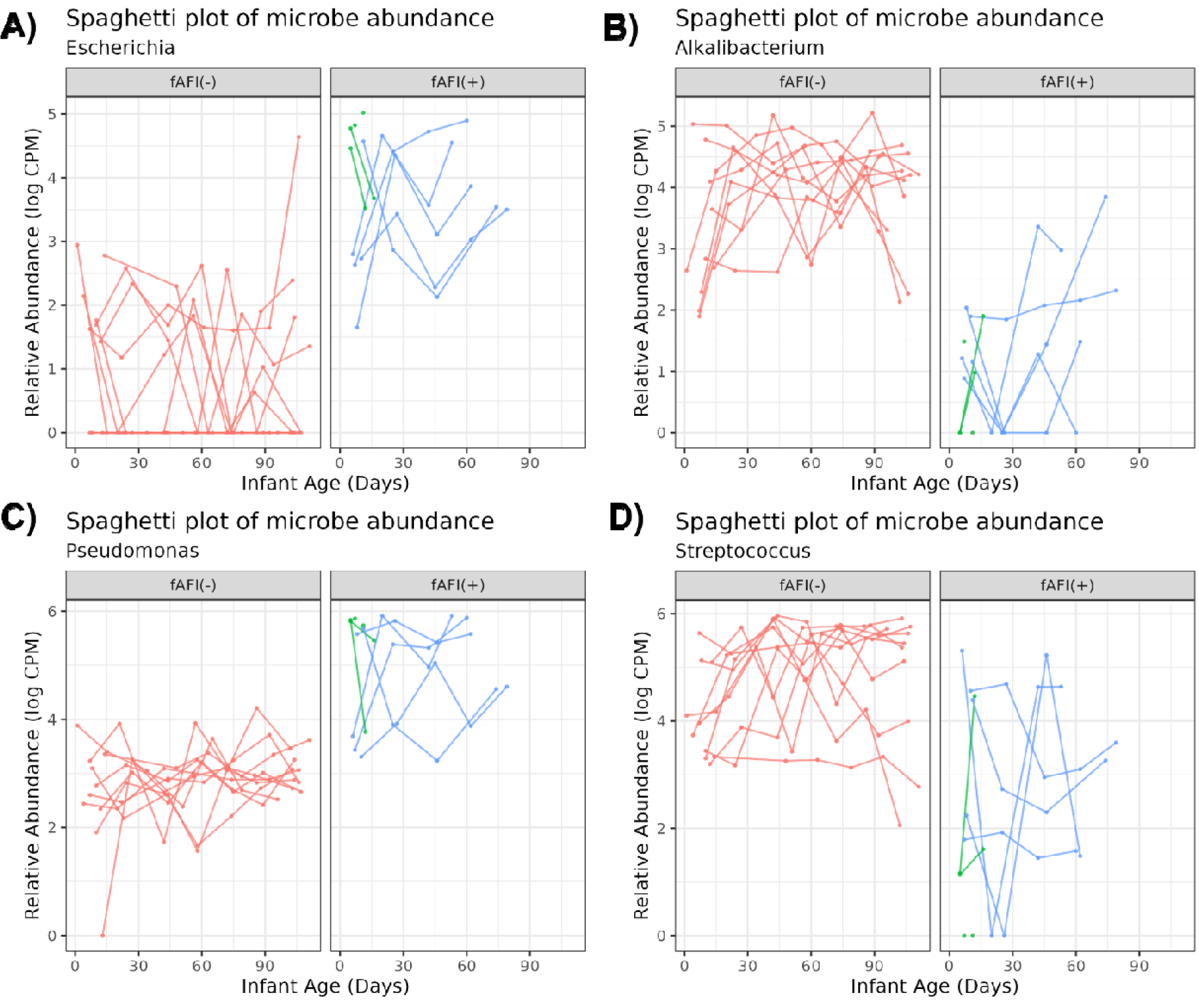
Spaghetti plots for the genera A) *Escherichia*, B) *Alkalibacterium*, C) *Pseudomonas*, and D) *Streptococcus*. These genera were chosen based on the visibly strong delineation between status of fatal acute febrile illness (fAFI) groups. fAFI(-) samples are shown in red, fAFI(+) early group samples are shown in green, and AFI(+) late group samples are shown in blue. In general, for fAFI(+) samples, *Escherichia* and *Pseudomonas* tended to be in higher abundance, whereas *Alkalibacterium* and *Streptococcus* tended to be in lower abundance.

### Beta Diversity

We next sought to analyze the pattern of overall community structure within fAFI groups. A uniform manifold approximation and projection analysis (UMAP) on the relative abundance of genera across fAFI statuses revealed that samples formed two distinct clusters based on their microbial profiles (Figure 4). In most cases, fAFI(+) infants congregated together in a tightly consolidated cluster, whereas fAFI(-) infants were widely divergent on the second dimension (Figure 4A). Furthermore, the fAFI(+) cluster was clearly separated from the fAFI(-) cluster. We also see that the fAFI(+) infants cluster is marked by *Pseudomonas, Leuconostoc*, and *Weisella* (Figure 4B). The fAFI(-) infants are subtly divided into subclusters as well and are mostly grounded in high *Staphylococcus, Streptococcus, Moraxella, Haemophilus*, and *Dolosigranulum*.

**Figure 4:**
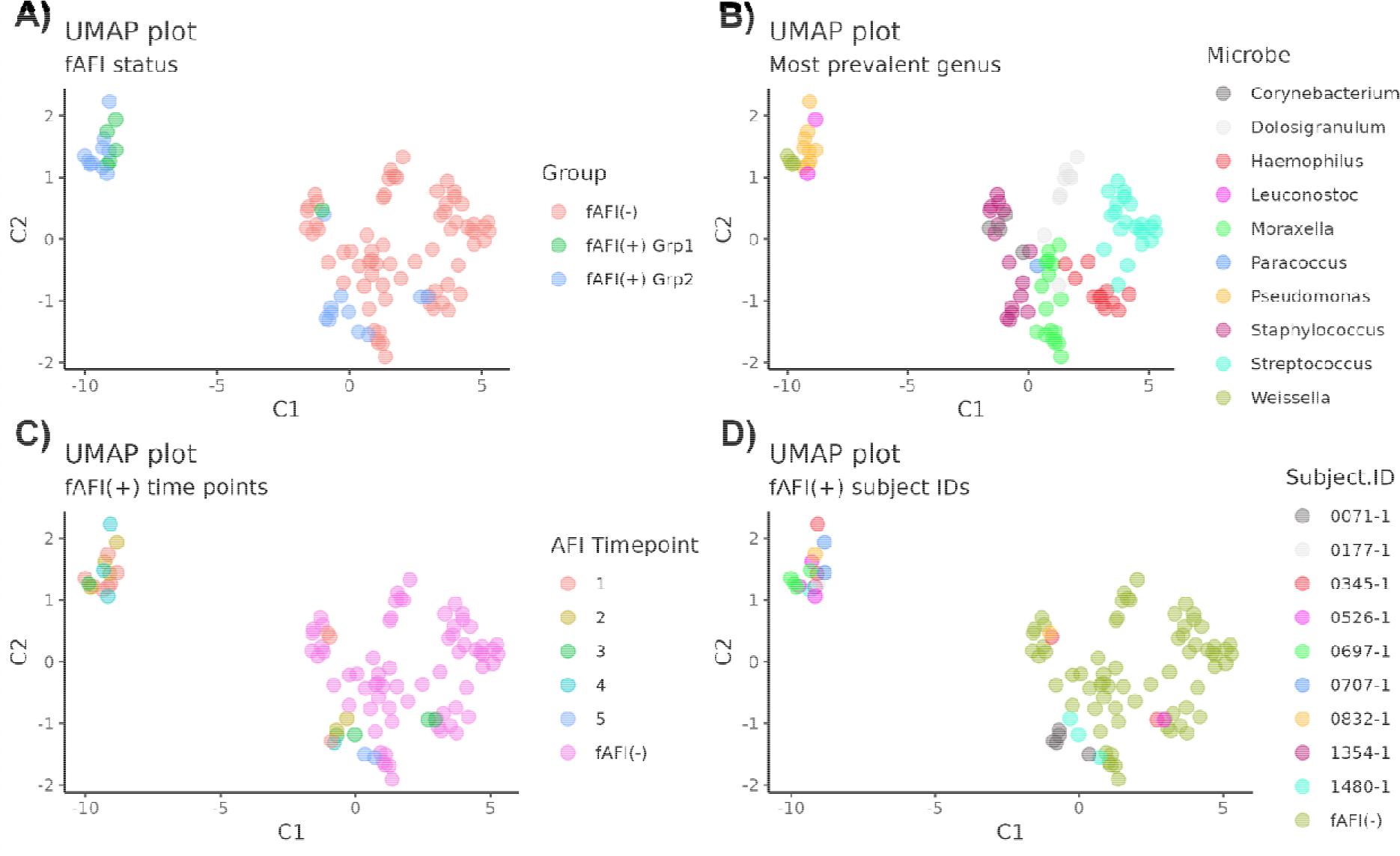
Panel of the same UMAP plot rendered with different colorations. The panels compare A) fatal acute febrile illness (fAFI) designations with subgroups, B) most prevalent genera, C) time point of sampling for fAFI(+) infants, and D) subject IDs for fAFI(+) infants. Strong delineations are visible between samples defined by certain genera and by fAFI status excepting samples belonging to a few subjects.

By highlighting the time point and subject ID (Figure 4C, 4D), some additional patterns emerge. Some of the samples from the late fAFI(+) group at early time points preceding fAFI cluster with the fAFI(-) group. Since this is prior to fAFI, it is not surprising to see similarities to the healthy fAFI(-). Second, the remaining fAFI(+) samples that cluster with the fAFI(-) samples belong to two infants, 1480-1 and 0071-1, and all of the samples for these two infants cluster together (Figure 4C).

Overall, the UMAP provides well-defined separation between the two groups on both dimensions and indicates intra-group microbiome profile differences.

We calculated the Bray-Curtis dissimilarity between samples to understand the community differentiation between groups at individual time points. Overall microbiome composition varied significantly between fAFI(+) infants and fAFI(-) infants. We produced a non-metric multidimensional scaling (NMDS) plot to visualize the Bray-Curtis dissimilarity. Colored ellipses encompass the samples within each fAFI status group, and star-shaped points indicate the calculated centroids. After fitting the variables of fAFI status and time point onto the ordination, time point was found to have an R^2^ of 0.27 and a p-value of 0.001, and fAFI status had an R^2^ of 0.4287 with a p-value of 0.001. Based on these values, the permutation goodness of fit test for both variables indicated that the groups are more related to the ordination axes of fAFI status and sample time point than expected by chance, indicating that the differences visible in microbial abundances are likely related to fAFI status and timepoint.

### Differential Abundance Analysis

To compare individual genus abundances directly between groups, we performed two sets of t-tests. First, we analyzed abundances between fAFI(+) and fAFI(-) infant groups (Table 2). When comparing the fAFI(-) to the early fAFI(+) group, we identified increased abundances of *Alkalibacterium* (adj. p=0.024), *Dolosigranulum* (adj. p=0.001), and *Moraxella* (adj. p=0.023) with decreased abundance of *Escherichia* (adj. p=0.053) in the AFI(-) infants.

**Table 2:**
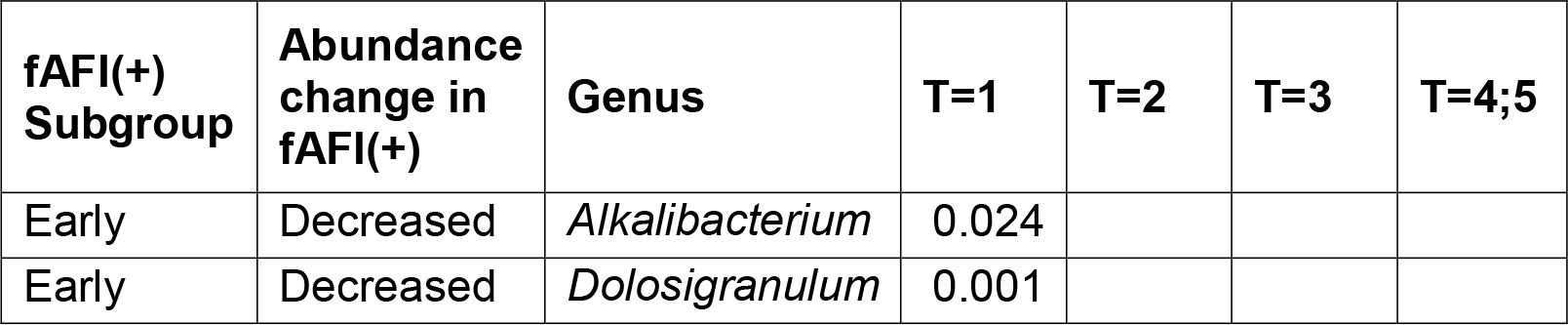
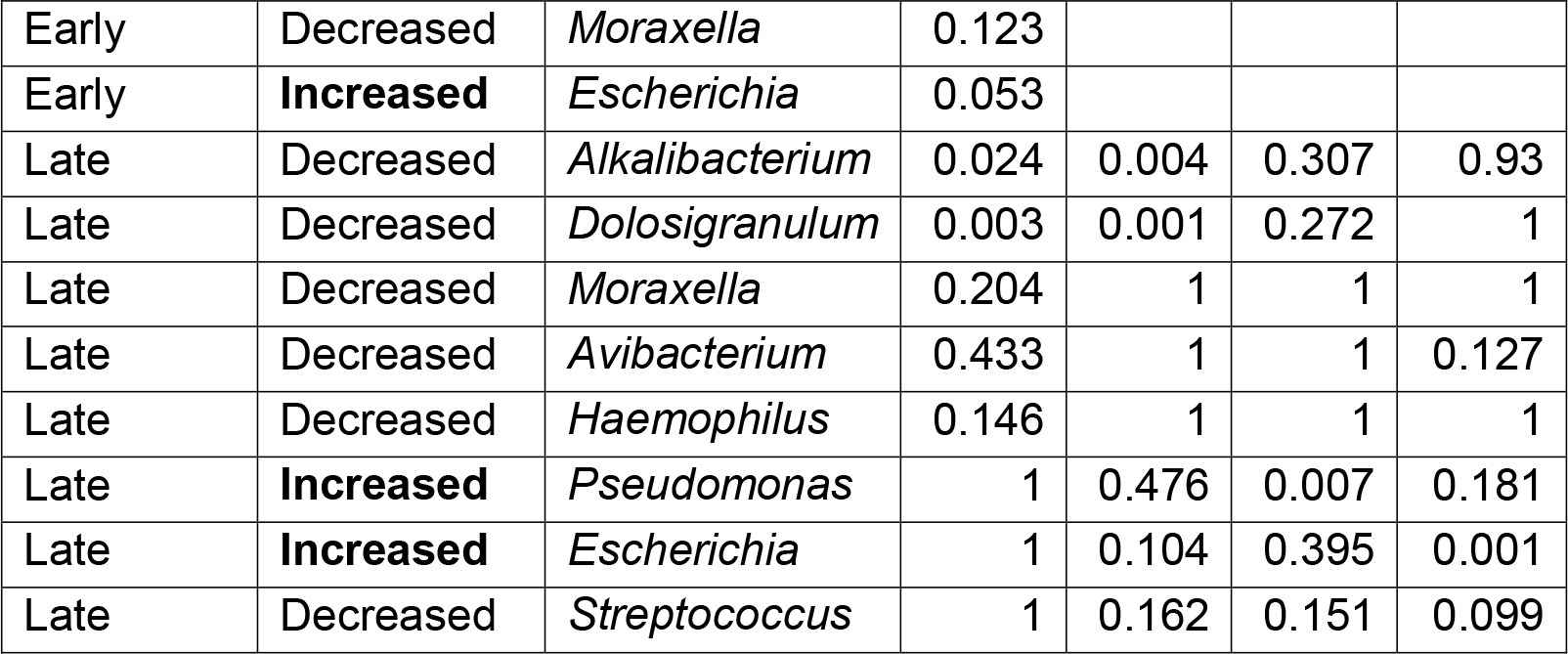
P-values from the comparison of fatal acute febrile illness (fAFI) status groups separated by early and late subgroups. The early subgroup was tested at the first time point, and the late subgroup was tested at time points 1, 2, 3 and combined at 4 and 5. Only genera with differences of import were included. Genera are marked as having increased or decreased in abundance relative to the control fAFI(-) group.

| fAFI(+) Subgroup | Abundance change in fAFI(+) | Genus | T=1 | T=2 | T=3 | T=4;5 |
| --- | --- | --- | --- | --- | --- | --- |
| Early | Decreased | *Alkalibacterium* | 0.024 |  |  |  |
| Early | Decreased | *Dolosigranulum* | 0.001 |  |  |  |
| Early | Decreased | *Moraxella* | 0.123 |  |  |  |
| Early | **Increased** | *Escherichia* | 0.053 |  |  |  |
| Late | Decreased | *Alkalibacterium* | 0.024 | 0.004 | 0.307 | 0.93 |
| Late | Decreased | *Dolosigranulum* | 0.003 | 0.001 | 0.272 | 1 |
| Late | Decreased | *Moraxella* | 0.204 | 1 | 1 | 1 |
| Late | Decreased | *Avibacterium* | 0.433 | 1 | 1 | 0.127 |
| Late | Decreased | *Haemophilus* | 0.146 | 1 | 1 | 1 |
| Late | **Increased** | *Pseudomonas* | 1 | 0.476 | 0.007 | 0.181 |
| Late | **Increased** | *Escherichia* | 1 | 0.104 | 0.395 | 0.001 |
| Late | Decreased | *Streptococcus* | 1 | 0.162 | 0.151 | 0.099 |

Patterns for these microbes were less defined across the various time points for the late fAFI(+) group (Table 2). *Alkalibacterium* and *Dolosigranulum* were both distinctly more present in the fAFI(-) group at time points 1 and 2 (adj. p<0.05) and differences became less clear closer to fAFI development (adj. p>0.05 at time points 3 and 4/5). Distinctions between groups are still evident closer to the development of fAFI in the late fAFI(+) subgroup. At the last sampling point, increases in *Escherichia* (adj. p=0.001) and *Pseudomonas* were present in the late fAFI(+) group (adj. p=0.181). This trend in *Pseudomonas* was also evident in the penultimate sampling (adj. p=0.007). *Streptococcus* was more abundant in the AFI(-) infants at the last sampling (adj. p=0.099).

We also analyzed the fAFI(+) profiles of infants to determine commonalities in microbial abundances at different time points of fAFI presentation. When t-tests were conducted between fAFI(+) subgroups at the sampling closest to AFI for individual genera, we found that their relative abundances were not significantly different (adj. p=1 for all tests), suggesting similar microbial profiles at time of fAFI.

## Discussion

The aim of this analysis was to investigate longitudinal changes in the nasopharyngeal microbiome of infants with fAFI as compared to healthy control infants. Based on our analysis, we believe that microbial identifiers distinguish fAFI(+) infants from their healthy counterparts. Our data suggest that the antecedent NP microbiome patterns of infants with fAFI were remarkably distinct from healthy infants and that dysbiosis precedes the onset of clinical disease by weeks, or even months. Whether this indicates that dysbiosis itself plays a causal or contributory role to the evolution of sepsis or is merely reflective of an underlying yet unmeasured mechanism (such as the infant’s underlying immunologic status) cannot be determined from our results.

The human nasopharynx is commonly occupied by a diverse array of microorganisms, most notably the bacterial residents *Moraxella, Staphylococcus, Corynebacterium, Dolosigranulum, Haemophilus*, and *Streptococcus*^*21,22*^. These microbes are found across timepoints in fAFI(-) infants, but dysbiosis proliferates in the fAFI(+) infants (**Figure 2**). Furthermore, in the upper respiratory tract, *Dolosigranulum* and *Corynebacterium* have been strongly associated with respiratory health as protective safeguards in the face of pathogens, notably *Streptococcus pneunominae*^*12,23-26*^. In our study, *Dolosigranulum* and *Corynebacterium* were almost nonexistent in the fAFI(+) infants (**Figure 2**) with fAFI(+) samples having a significantly lower abundance of *Dolosigranulum* (p<0.003, Table 2). This may reduce the protective effect of these microbes in fAFI(+) infants and may cause increased susceptibility to disease.

In the context of fAFI, several genera were of interest across our performed analyses and visualizations; specifically, we saw higher abundances of *Escherichia, Pseudomonas, Leuconostoc*, and *Weissella* in fAFI(+) infants. This pattern is visible in the stacked bar and spaghetti plots (Figures 2, 3A, 3C) and confirmed for the latter three genera in the UMAP plot renderings (Figure 4A, 4B). T-tests also confirmed significantly higher abundances of *Escherichia* and *Pseudomonas* at the times close to fAFI occurrence. *Escherichia* is often found in the gut of healthy infants but is known to cause a number of diseases when found elsewhere, including diarrhea and pneumonia^27-30^. *Pseudomonas* has been implicated as a successful upper airway colonizer in patients with lung damage^31,32^ and efforts to control *Pseudomonas* reservoirs in a hospital setting significantly reduced rates of sepsis and pneumonia in autopsied infants^33^. While no direct causative links between either *Escherichia* or *Pseudomonas* have been found with AFI, their presence is abnormal in healthy microbiota. On the other hand, *Leuconostoc* and *Weissella* have been identified in breast milk^34^ and feces^35^ and have been associated with environmental sources of potential contamination^36-38^. Due to the separate sequencing of controls and fAFI samples, we cannot definitively excuse *Leuconostoc* and *Weisella* from being environmentally linked contaminants.

Across all sepsis time points, there was generally lower beta diversity in the fAFI(+) samples, and the strength of the inter-sample differences was verified with the NMDS goodness of fit tests (**Figure 5**). Lower diversity in the NP microbiome has been previously associated with increased occurrence of rhinovirus illness^39^ and HIV-associated bronchiectasis^40^, implying that greater NP diversity signals a healthy microbiome^41^.

**Figure 5:**
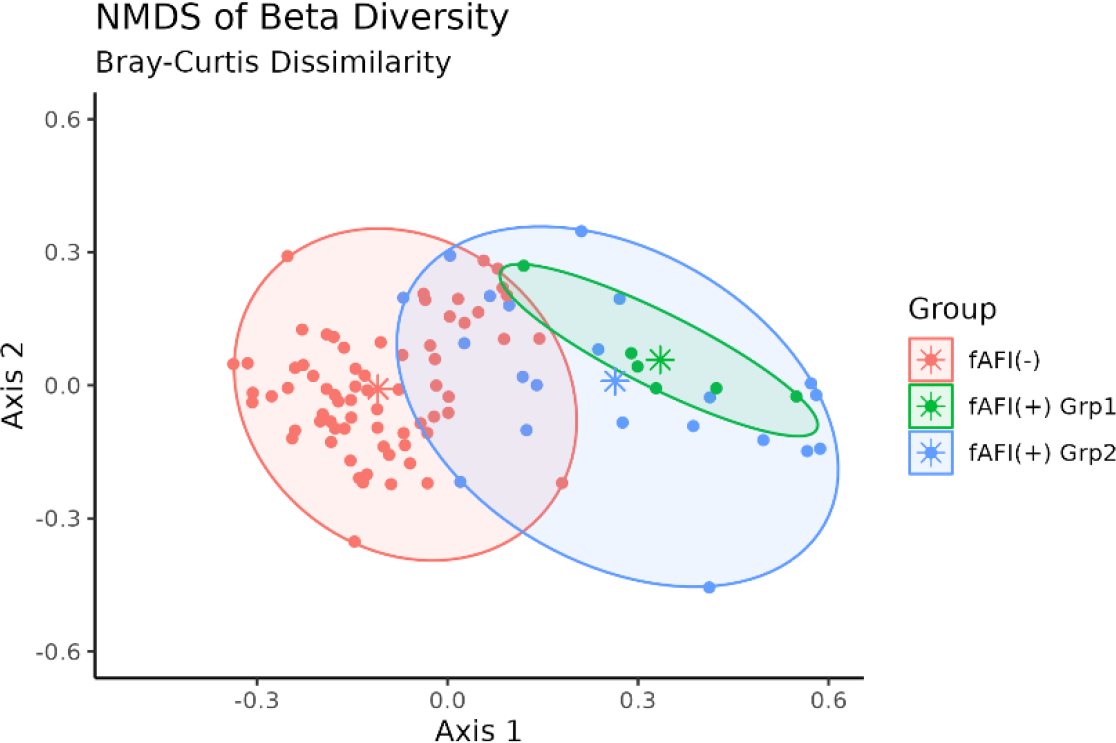
A non-metric multidimensional scaling (NMDS) plot for fatal acute febrile illness (fAFI) status and sample timepoint of the calculated Bray-Curtis dissimilarity. Dissimilarity was calculated as a measure of beta diversity. The stress value was 0.2. Ellipses colored by fAFI status and group encircle the dissimilarity values in nondimensional space. Group centroids are demarcated by a star shaped point in the center of each ellipse. Samples have clustered based on fAFI status and timepoint, indicating that there are observable differences in the microbiome of the fAFI(-) and fAFI(+) groups.

One limitation of our study is the small number of infants included in the study. Overall, we have *N*=96 observations, yet the number of fAFI(+)/-subjects in these analyses are 9 and 10, respectively, and thus the dataset results in a small number of points when separated by time of fAFI. This results in limited options for statistical comparison and limits specific conclusions at each time point. For example, we found that the differential abundance differences between fAFI(+) and fAFI(-) infants were most prevalent at the timepoints directly before an fAFI event. However, these differences might be due to drops in fAFI(+) sample number caused by infant drop out due to death shortly after fAFI, leading to discernable differences prior to the occurrence of fAFI. The sample size is large enough to make generalized conclusions across all time points and provide hypotheses for individual time points for future study. We therefore suggest that further study and validation of our findings be conducted.

In conclusion, we have found that microbiome dysbiosis precedes fAFI, in some cases before the onset of initial infection. We recommend that further studies be carried out to validate our results and further investigate NP microbiome differences in the infants with fAFI, specifically with a larger longitudinal study. We also recommend investigating the interaction between NP and gastrointestinal microbiomes in sick infants because we believe that dysregulation is likely not limited to the nasopharynx. Lastly, we note a need to investigate the mechanisms of action of distinctive microbes to better elucidate their role in disease acquisition.

## Supporting information

Supplementary data processing and sequencing methods

## Data and Code

Data available at NCBI BioProject accession number PRJNA1021270. https://identifiers.org/NCBI/BioProject:PRJNA1021270

Source code available from: https://github.com/aubreyodom/FAFI-microbiome Archived source code at time of publication: DOI: 10.5281/zenodo.8380998

## Author Contributions

The study was conceived by C.G., R.L., and W.E.J. Data were curated by W.M., R.L. and A.R.O. Formal data analysis was conducted by A.R.O., and J.M. Methodology was conspired by A.R.O., J.M., R.L., W.M., and W.E.J. Project administration was carried out by R.P. and A.R.O. Resources were provided by A.I. Software, validation, visualization, and original draft preparation were carried out by A.O. Supervision was conducted by W.E.J., C.G., and R.L. All authors reviewed the manuscript prior to submission and provided feedback.

