## Supplementary data processing and sequencing methods for "Analysis of nasopharyngeal microbiome patterns in Zambian infants with fatal acute febrile illness"

**Supplementary Material**

Sample Processing Methods

*Sample processing and storage*

NP swabs were obtained from the posterior nasopharynx using a sterile flocked tipped nylon swab (Copan Diagnostics, Murrieta, California). The swabs were then placed in universal transport media, put on ice and transferred to our onsite lab on the same campus, where they were aliquoted and stored at -80°C until DNA extraction. DNA was extracted using the NucliSENS EasyMagG System (bioMérieux, Marcy l’Etoile, France). Extracted DNA was stored at a lab located at the University Teaching Hospital in Lusaka at -80°C. Sample collection, processing and storage were previously described (Gill et al., 2016).^1^

*16S ribosomal DNA amplification and MiSeq sequencing*

Library preparation was performed according to the standard instructions of the 16S Metagenomic Sequencing Library preparation protocol (Illumina, Inc., San Diego, CA, United States). Amplicons were indexed using IDT for Illumina Nextera DNA Unique Dual Indexes (Illumina Inc., San Diego, CA, USA) according to the manufacturer’s recommendation. The amplicon library concentration was measured on the Qubit® 3.0 Fluorometer (Invitrogen) and the size of the amplicons were then visualised using the 4200 TapeStation (Agilent Technologies, Santa Clara, CA, USA). A negative control was used in the amplicon preparation step. No amplification was observed in the negative control. The purified amplicon libraries were normalized to 2 nM and then sequenced on the Illumina MiSeq platform using the Illumina MiSeq v3 kit (Illumina Inc.), to obtain 2 x300 bp paired-end sequences. Sequencing was performed at the Sequencing Core Facility, National Institute for Communicable Diseases, South Africa. Each group was sequenced at separate times as part of separate manifests. Lab technicians were blinded to the timing of sample collection and clinical data.

*Quality control and annotation of isolates*

The quality of sampling of the NP swabs was confirmed initially using PCR against the constitutive human enzyme RNAseP. All samples included in this analysis were RNAseP positive, which demonstrates adequacy of the sample collection procedure in terms of making contact with the respiratory mucosa, and the absence of endogenous PCR inhibitors.

We assessed the quality of the sequencing data using FastQC v0.11.7^2^ and compiled reports using MultiQC v1.10.1. Trimmomatic v0.39^3^ was used to trim Illumina adapters and remove low-quality sequences. We performed a sliding window trim, cutting once when the average quality score within a window of four bases falls below 20. We removed both leading and trailing low quality or N bases below quality three. All reads were trimmed with a head crop at 30 bases and crop at 225 bases. All other parameters used the default settings. Quality control reports are hosted on the Zenodo repository of code for this analysis.

1.	Gill CJ, Mwananyanda L, MacLeod W, et al. Incidence of Severe and Nonsevere Pertussis Among HIV-Exposed and -Unexposed Zambian Infants Through 14 Weeks of Age: Results From the Southern Africa Mother Infant Pertussis Study (SAMIPS), a Longitudinal Birth Cohort Study. *Clin Infect Dis*. Dec 01 2016;63(suppl 4):S154-S164. doi:10.1093/cid/ciw526

2.	Andrews S. FastQC: a quality control tool for high throughput sequence data. Babraham Bioinformatics, Babraham Institute, Cambridge, United Kingdom; 2010.

3.	Bolger AM, Lohse M, Usadel B. Trimmomatic: a flexible trimmer for Illumina sequence data. *Bioinformatics*. 2014;30(15):2114-2120.
